# *Pantr2*, a trans-acting lncRNA, modulates the differentiation potential of neural progenitors in vivo

**DOI:** 10.1101/2022.10.07.511381

**Authors:** Jonathan J Augustin, Saki Takayangi, Thanh Hoang, Briana Winer, Seth Blackshaw, Loyal A Goff

## Abstract

Ablation of the long non-coding RNA (lncRNA) *Pantr2* results in microcephaly in a knockout murine model of corticogenesis, however, the precise mechanisms used are unknown. We present evidence that *Pantr2* is a trans-acting lncRNA that regulates gene expression and chromatin accessibility both in vivo and in vitro. We demonstrate that ectopic expression of *Pantr2* in a neuroblastoma cell line alters gene expression under differentiating conditions, and that both loss and gain of function of *Pantr2* results in changes to cell-cycle dynamics. We show that expression of both the transcription factor *Nfix* and the cell cycle regulator *Rgcc* are negatively regulated by *Pantr2*. Using RNA binding protein motif analysis and existing CLIP-seq data, we annotate potential HuR and QKI binding sites on *Pantr2*, and demonstrate that HuR does not directly bind *Pantr2* using RNA immunoprecipitation assay. Finally, using Gene Ontology enrichment analysis, we identify disruption of both Notch and Wnt signaling following loss of *Pantr2* expression, indicating potential *Pantr2*-dependent regulation of these pathways.

## Introduction

The mammalian cerebral cortex is a highly ordered and evolutionarily conserved tissue in the central nervous system. The cortex is organized into six molecularly distinct laminae (I-VI) (Molyneaux et al., 2007). Proper cortical development requires a complex orchestration of specific processes, both cell-intrinsic and cell-extrinsic, that coordinate temporally and spatially to give rise to the mature cortex (Cadwell et al., 2019). Interestingly, the cortex expresses more long noncoding RNAs (lncRNAs) than any other somatic tissue, raising questions about their potential involvement in establishing the diversity of cortical neurons and their organization (Cabili et al., 2011; Mercer et al., 2008). lncRNA transcripts are 200nt in length with no predicted protein-coding potential. Roughly 40 % of annotated lncRNAs are uniquely expressed within the CNS, many of which are dynamically expressed and spatiotemporally restricted (Belgard et al., 2011; Cabili et al., 2011; D. Li & Yang, 2017; Molyneaux et al., 2007; Molyneaux et al., 2015; Sauvageau et al., 2013). This number is likely an underestimation, as it does not reflect the abundance of unannotated lncRNAs commonly found to be specific to the CNS (D. Li & Yang, 2017). Current evidence suggests that lncRNAs can act as epigenetic regulators of gene expression, regulating splicing, transcription, translation, and mRNA degradation (Perry & Ulitsky, 2016; Rinn & Chang, 2012). lncRNAs such as Dali, Pnky, and NORAD have been attributed roles in regulating corticogenesis, suggesting that other lncRNAs may also play important regulatory roles in this process (Andersen et al., 2019; Chalei et al., 2014; Elguindy & Mendell, 2021). Developing a clear understanding of the roles and molecular mechanisms lncRNAs employ to regulate corticogenesis is important in increasing our understanding of this highly complex process.

Previous work has identified a novel lncRNA – *Pou3f3* adjacent non-coding transcript 2 (*Pantr2*) – as necessary for the proper development of the neocortex (Goff et al., 2015; Lai et al., 2015; Sauvageau et al., 2013). *Pantr2* is 462nt in length and is expressed in embryonic neural stem cells as well as the adult brain and kidney in mice. Although the *Pantr2* locus is 8kb downstream of the *Pou3f3* locus and many lncRNAs have cisregulatory effects on neighboring genes, it is unclear whether *Pantr2* exerts a biological effect through a *Pou3f3*-dependent pathway. Expression of *Pantr2* in the developing cortex has been observed as early as embryonic day 13.5 (E13.5) in the lateral and medial ganglionic eminence (Goff et al., 2015; Sauvageau et al., 2013). During cortical development, this expression expands to the ventricular and subventricular zone before being postnatally restricted to the somatosensory cortex, visual cortex, and the olfactory bulb. Deletion of the *Pantr2* gene locus in a murine model leads to an expansion of deep layer cortical projection neurons and a reduction in upper layers, resulting in an overall microcephalic phenotype (Goff et al., 2015; Sauvageau et al., 2013). This is accompanied by a decreased number of intermediate progenitor cells (IPCs) generated in the neocortex of *Pantr2*^-/-^ animals relative to wild-type littermates, with no significant difference detected in the number of apical progenitor cells (APCs). However, the mechanisms through which *Pantr2* deletion alters corticogenesis are unknown. Here, we provide insights into the molecular mechanisms used by *Pantr2* to direct neocortical development and fate specification. We also identify key downstream effector genes impacted by *Pantr2* expression and likely contribute to the observed phenotypes within this murine knockout model.

We demonstrate that the *Pantr2* lncRNA is a trans-acting regulator of the cell cycle in neural progenitor cells (NPCs) and that this activity is specific to contexts in which NPCs are actively committing to a differentiated neuronal state. Using single-cell genomic assays, we identify changes in the expression of genes associated with cell cycle and Wnt signaling consistent with a microcephalic phenotype in the *Pantr2* knockout. We identify changes in the expression of the transcription factor nuclear factor I X (*Nfix*) which regulates the proliferation and migration of cells in the subventricular zone (SVZ), a major neurogenic niche located in the developing cortex (Heng et al., 2014; Mason et al., 2009; Zalucki et al., 2019). Additionally, we identify Regulator Gene of Cell-Cycle (*Rgcc*) as a potential novel downstream effector of Pantr2 involved of fate specification in the cortex. Expression of *Rgcc* is linked with stalled progression through mitosis, and thus longer time spent in the cell cycle (Saigusa et al., 2007). Finally, publicly available CLIP-seq data indicates that the RNA binding protein quaking (Qki) binds to mature *Pantr2* transcripts in embryonic day 14.5 (E14.5) dorsal telencephalon, presenting a new molecular target for the *Pantr2* RNA.

## Methods

### Mouse generation, processing, and housing

Wildtype and *Pantr2* knockout mice were maintained on a C57BL/6J background. *Pantr2* knockout mice were obtained from the Rinn Lab and were generated using the VelociGene method described in Sauvageau et al. 2013. Timed pregnancies were arranged by mating breeders in the evening and separating them the following morning. Dams with a plug were considered to have embryos at day 0.5 (E0.5). Embryos were allowed to develop for the desired number of days before the dam was euthanized by cervical dislocation and the embryos were surgically removed. The developing brain of each pup was dissected and placed in 4 % paraformaldehyde (PFA) for two hours at 4°C and moved into 30 % sucrose overnight. Brains were then embedded in optimal cutting temperature (O.C.T) media and stored at −80°C. All cryosections were done at 25 µm thickness, adhered to charged slides, and stored at −80°C for future use. Brains were dissected after perfusion and fixed in 4 % PFA for 2 hours at 4°C before being moved into 30 % sucrose overnight. Postnatal brains were embedded, sectioned, and stored in the same manner described for embryonic brains. All mice were housed in the Johns Hopkins School of Medicine animal care facilities in compliance with the Animal Welfare Act regulations and Public Health Service Policy.

### Generation of Neurospheres

The dorsal telencephalon of E14.5 knockout and wildtype mice were isolated through dissection and placed in 1ml HBSS solution. A P1000 tip was used to triturate the tissue gently and the resultant cell suspension was centrifuged at 500G for 5 minutes to pellet the cells. To these cell pellets, 7mL of Neurosphere Growth Media (480ml DMEM/F-12 with glutamine [cat: 11320-033], 1.45g Glucose, 2.5ml of 5,000 U/mL penicillin/streptomycin, 1X N2 supplement [cat: 17502048], 1X B-27 supplement without retinoic acid [cat: 12587010]) was added and the cells were resuspended and passed through a 40 µm cell strainer. These single cells were plated in ultra-low adherence T25 flasks and cultured until spheroids formed. These spheroids were passaged by resuspending them in 0.05 % TrypLE and incubating at 37°C for 5-10 minutes. Gentle trituration was used to generate a single cell suspension which was passed through a 40 µm cell strainer to remove large cell clumps. Neurospheres were stored for longterm use by slow freezing in Neurosphere Growth Media supplemented with 10 % DMSO at −80°C and subsequent storage in the liquid phase of liquid nitrogen.

### Neurosphere proliferation assay

Neurospheres were dissociated as described above and plated in ultra-low attachment 6 well plates at 10,000 cells per well using biological triplicates for *Pantr2*^-/-^ and wildtype neurospheres. Cells were cultured in Neurosphere Growth Media at 37°C and 5 % CO2. Every 24 hours for 7 days, images were taken at 4X magnification using 3 fields per well. Images were processed using a pipeline created in Cellprofiler (Stirling et al., 2021) to assess the area of objects within a given field for a total of 3 fields per replicate and 3 replicates per clone. Documentation of data analysis is provided in the GitHub repository associated with this manuscript (https://www.github.com/jaugust7/pantr2_mechanism_manuscript).

### Single-cell library preparation and data analysis for cortical neurospheres

Neurospheres were digested using 1X TrypLE and gently triturated. Neurospheres were then incubated at 37°C for 5 minutes before being centrifuged at 500G for 5 minutes. The TrypLE Enzyme was removed and the cells were resuspended in 1X PBS and passed through a 40 µm cell strainer. This single cell suspension was then processed using the 10X Genomics Chromium Single Cell V2 kit following the manufacturer’s directions and targeting 10,000 cells per sample. Single-cell libraries were uniquely indexed, pooled, and sequenced to an average depth of 50 million paired-end reads per sample on the Illumina NovaSeq 6000 platform. Reads were aligned against a mouse reference transcriptome (Gencode, vGRm38) (Frankish et al., 2019) using 10X Genomics Cell Ranger 3.1.0.

For each sample, cells with greater than 40,000 and less than 1,000 UMIs were discarded resulting in a final dataset of 25,343 cells. These cells were then used as the input into the Monocle3 single cell analysis framework. Documentation of data analysis is provided in the GitHub repository associated with this manuscript (https://www.github.com/jaugust7/pantr2_mechanism_manuscript).

### Cell culture conditions and N2a differentiation conditions

Neuro-2a (N2a) cells were cultured in high glucose DMEM with added 10 % fetal bovine serum (FBS) and 1 % Pen/Strep (Complete Growth Media) in tissue-culture treated dishes at 37°C and 5 % CO2. N2a differentiation was induced by removing the complete growth medium and replacing it with high glucose DMEM with added 1 % Pen/Strep (Serum Free Media). Cells were subjected to this differentiation protocol for 48 hours before collection and analysis. N2a cells were transfected using FuGENE HD following the manufacturer’s specifications with a DNA to lipofection reagent ratio of 1:5. Cells were harvested 48 hours after transfection for RNA isolation and western blot analysis.

### Comparative gene expression analysis using qPCR

For comparative gene expression analysis in the N2a cells, we isolated RNA from cultured cells using the Qiagen RNeasy Mini Kit [cat: 74104] following the manufacturer’s instructions. This RNA was converted to cDNA using the SuperScript IV First-Strand Synthesis System [cat: 18091050] following manufacturer’s instructions. Finally, qPCR reactions were set up using ABI PowerUp SYBR Green Master Mix [cat: A25742] following the included instructions using primers indicated in **Supplemental Table 1**.

### Single molecule fluorescence in-situ hybridization and immunofluorescence

Single-molecule fluorescence in-situ hybridization (smFISH) was performed using hybridization chain reaction (HCR) (Choi et al., 2018). A probe set was designed against *Pantr2* using the Python package HCRProbeDesign available at the Github repository (https://www.github.com/gofflab/HCRProbeDesign). Briefly, E13.5 whole heads were fixed in 4 % PFA overnight before being moved into 30 % sucrose until the tissue equilibrated with the sucrose solution. The heads were then embedded in O.C.T media on a dry ice and ethanol slurry before being stored at −80°C. These embedded tissues were sectioned at a thickness of 25 µm adhered to charged slides, and stored at −80°C until ready for HCR. Sections were removed from −80°C and allowed to warm to room temperature. Slides were then immersed in 50 % EtOH, 70 % EtOH, and 100 % EtOH for 5 minutes each at room temperature. After a final 5 minute wash in 100 % EtOH, the sections were washed twice in 1X PBS for 5 minutes each. The 30 % probe hybridization buffer (30 % formamide, 5X SSC media, 9mM citric acid, 0.1 % Tween-20, 100 µg/mL heparin, 1X Denhardt’s solution, and 10 % dextran sulfate) was pre-warmed to 37°C. Slides were immersed in 30 % probe hybridization buffer and allowed to pre-hybridize for 10 minutes at 37°C in a humidified chamber. The probe solution was prepared by diluting 0.4 µL of 1 µM stock HCR oligos with 100 µL of 30 % probe hybridization buffer. The prehybridization buffer was removed and replaced with 150 µL of the probe solution and incubated overnight at 37°C in a humidified chamber.

After incubation, the probe solution was removed and slides were washed in 30 % probe wash buffer (30 % formamide, 5X SSC media, 9mM citric acid, 0.1 % Tween-20, 100 µgml heparin) at 37°C. Next, slides were washed in 75 % 30 % probe wash buffer 25 % 5X SSC-Tween-20 for 15 minutes, 50 % 30 % probe wash buffer 50 % 5X SSC-Tween-20 for 15 minutes, 25 % 30 % probe wash buffer 75 % 5X SSC-Tween-20, and finally 100 % 5X SSC-Tween-20 at 37°C (5X SSC-Tween-20 contains 0.1 % Tween-20). Slide edges were dried and then 200 µL of amplification buffer (5X SSC buffer, 0.1 % Tween-20 was added to each slide and incubated at 37°C for 30 minutes. During this incubation period 6pmol of both hairpin H1 and hairpin H2 (Molecular Instruments B2-647) were prepared separately in 100 µL of amplification buffer per slide. The hairpins were then snap cooled by heating them to 95°C for 90 seconds and cooling in the dark for 30 minutes at room temperature. Hairpins were added to each other 1:1 after snap cooling. After the 30 minute incubation in amplification buffer, slides were drained of amplification buffer and 150 µL of the combined hairpins were added to each slide and incubated overnight at 37°C. After this incubation, excess hairpins were removed by incubating in 5X SSC-Tween-20 at room temperature twice for 30 minutes before applying counterstaining in DAPI and sealing the slides for imaging.

### Single nuclei sequencing library preparation and data analysis for dorsal telencephalon

Day E15.5 embryonic brains were collected from both wildtype and *Pantr2*^-/-^ mice. The dorsal telencephalon was dissected, flash frozen in liquid nitrogen, and stored at −80°C until ready for nuclei isolation. Nuclei were isolated using a modified version of the Salty-EZ Lysis Buffer method outlined on protocols.io (https://tinyurl.com/36ryzay4). In brief, tissues were lysed in 300 µL of ice cold Salty-EZ10 lysis buffer containing 10mM Tris HCL pH 7.5, 146mM NaCl, 1mM CaCl2, 21mM MgCl_2_, 0.03 % Tween-20, 10 % EZ-lysis Buffer [Sigma Aldrich], 1mM DTT and 0.2 µLl Protector RNase Inhibitor. The tissue was gently triturated using a P200 wide-bore pipette tip on ice until the tissue was completely in solution. Once homogenized, 700 µL of ice cold Slaty-EZ10 buffer was added and each sample was gently mixed using a wide-bore pipette tip. Next, the solution was spun down at 4°C at 500G for 5 minutes to pellet the nuclei in suspension. The supernatant was aspirated off and 500 µL of WRB2 buffer containing 10mM Tris-HCl pH 7.5, 10mM NaCl, 3mM MgCl_2_,1 % w/v bovine serum albumin (BSA), 1mM DTT and 0.2 µL Protector RNase Inhibitor. Nuclei were washed three times in 500 µL WRB2 buffer before being used as the input material for 10X Genomics Chromium Single Cell V2 kit following standard protocol included with the kit.

Libraries were sequenced using an Illumina NovaSeq to approximately 50 million reads per sample on an S1 flowcell. After sequencing, reads were aligned using the Kalistobus pipeline against GRCm38 index generating both spliced and unspliced read matrices. These matrices were combined and these counts were used as input into Diem, an R package designed to identify a signature for debris droplets and identify test droplets with enrichment for these debris signatures (Alvarez et al., 2020). Post debris cleanup, the data was subjected to doublet removal using the R package scds (Bais & Kostka, 2020) and manual identification and removal of cells of undesired origin including the ventral telencephalon, microglia, fibroblast and ependymal cells based on marker gene expression resulting in a final dataset of 67K nuclei split evenly between samples. Documentation of data analysis is provided in the GitHub repository associated with this manuscript (https://www.github.com/jaugust7/pantr2_mechanism_manuscript).

### ATAC-seq library preparation and data analysis

Neurospheres were passaged using 1X TrypLE Express [Thermofisher cat: 12604013] with gentle trituration and passed through a 40 µm cell strainer. For ATAC-seq library preparation, we utilized the Omni-ATAC-seq protocol first described by (Buenrostro et al., 2013; Buenrostro et al., 2015), with minor changes. For each biological replicate, 50,000 cells were counted and washed in ice-cold 1ml 1X PBS. To isolate nuclei, the cells were pelleted and resuspended in 50 µL of ice cold Lysis Buffer (10mM Tris-HCl, pH 7.5, 10mM NaCl, 3mM MgCl_2_, 0.1 % NP-40, 0.1 % Tween-20,0.01 % Digitonin) and gently mixed by pipet 3 times to resuspend. Cells were incubated on ice in Lysis Buffer for 3 minutes before being washed with 1ml Wash Buffer (10mM Tris-HCl, pH 7.5, 10mM NaCl, 3mM MgCl_2_, 0.1 Tween-20) 3 times to stop the lysis reaction. Nuclei were spun down at 500G for 10 minutes and the supernatant was discarded. For each sample, a Transposition Reaction Mix was made using the Nextera DNA Library Prep Kit (25 µL 2X TD Buffer (Tagment DNA Buffer), 16.5 µL 1X PBS, 0.5 µL 10 % Tween-20, 0.5 µLl 1 % Digitonin, 2.5 µL Tn5 Transposase (Tagment DNA Enzyme 1), 5 µL nuclease-free H_2_O. Each nuclear pellet was resuspended in the Transposition Reaction Mix and incubated at 37°C for 30 minutes in a heat block with occasional mixing. DNA was purified using a Qiagen MinElute Reaction Cleanup Kit and eluted in 10 µL of 10mM Tris-HCl pH 8.0. The isolated DNA was used as the input into the PCR amplification/Library Generation step. Each PCR reaction consisted of 10 µL of purified transposed DNA, 10 µL nuclease-free H_2_O, 2.5 µL of Ad1_noMx forward primer, and 2.5 µL of the reverse primer associated with the sample **(Supplemental Table 2)**. Samples we PCR amplified using the following settings: 72°C for 5 minutes, then 98°C for 30 seconds followed by 16 cycles of (98°C for 10 seconds, 63°C for 30 seconds, 72°C for 1 minute) then 72°C for 5 minutes and hold at 4°C. The number of PCR cycles to use was determined by using qPCR on 1/5th of the library from each sample. After the first 5 cycles above, the libraries were removed and 5 µL of the PCR reaction was subjected to qPCR for 20 cycles. The number of cycles needed to reach a third of the maximum fluorescence for each sample was used as the remaining number of cycles to add, in this case, 11 cycles, for a total of 16 cycles.

Libraries were purified using AMPure XP beads to remove primer dimers using the standard protocol. To remove the small fragments, 1.8X bead volume was added to each library. For removing large fragments >1000bp, 0.5X bead volume was used on the eluate obtained above. Once finished, the purified libraries were quality controlled using Agilent High Sensitivity DNA bioanalysis chip to look for nucleosome periodicity and measured using QuBit 2.0 for DNA concentration, both following manufacturer’s instructions. Samples were sequenced using 50bp paired-end reading at roughly 50 million reads per sample.

Sequenced libraries were aligned to the mouse genome (UCSC mm10) using the bowtie2 short read aligner. From here, bam files were generated and indexed using samtools, and bigwig files were generated using bamCoverage. Peaks were called using MACS-3 and differential peak analysis was performed in R using DiffBind. Motif enrichment analysis was performed using the R implementation of MEME Suite’s Analysis of Motif Enrichment test. Documentation of data analysis is provided in the GitHub repository associated with this manuscript (https://www.github.com/jaugust7/pantr2_mechanism_manuscript).

### Thymidine analog incorporation assay and flow cytometry analysis

Thymidine analog incorporation was accomplished using the Invitrogen EdU Click-iT kit [ThermoFisher Scientific cat: C10425] following the manufacturer’s directions. Samples were subjected to flow cytometry using linear scaling for the DNA content channel (DAPI) and log scale for the EdU incorporation channel (Alexa Fluor 488). Flow cytometry results were imported into Floreada.io, a free, web-based tool for simple flow cytometry analysis. The gating strategy used is presented in **(Supplemental Figures 1A and 1B)**.

### Site-directed mutagenesis

Site-directed mutagenesis for construction of the *Pantr2* deletion mapping experiment was performed using the Q5 Site-Directed Mutagenesis Kit from New England Biosciences following the manufacturers’ protocol [cat: E0554S]. The primers used for each deletion mutant are listed in **(Supplemental Table 3)**.

### Native RNA immunoprecipitation and qPCR

N2a cells were grown in 10cm dishes in 5 % CO2 at 37°C to 90 % confluence. Cells were collected using a cell lifter and washed twice in ice-cold PBS. N2a cells were then lysed using 100 µL ice cold PLB buffer (100mM KCl, 5mM MgCl_2_, 10mM HEPES pH 7, 0.5 % NP-40, 1mM DTT, 1X EDTA-free Protease inhibitor cocktail, 200U/mL RNase inhibitor) per 2 million cells and incubated on ice for 5 minutes before being incubated at −80ºC for at least 1 hour. Magnetic Protein A beads were used for immunoprecipitation (IP) of HuR protein and were prepared by washing 75 µL of resuspended beads, for each IP, twice with 500 µL of NT-2 buffer (50mM Tris-HCl ph 7.4, 150mM NaCl, 1mM MgCl_2_ and 0.05 % NP-40) using a magnetic rack to pellet the beads in between washes. The Protein A beads were then resuspended in 100 µL of NT-2 buffer and 5 µg of either HuR antibody [Invitrogen cat: 39-0600] for the HuR RIP or Mouse IgG [Diagenode cat: C15400001-15] for the IgG control RIP was added to experimental and control tubes in triplicate. The beads were incubated with antibodies for 1 hour at room temperature with rotation. These antibody-coated beads were washed six times with 1ml NT-2 buffer with gentle mixing between washes. After the washes, the beads were resuspended in 900 µL of NET-2 buffer (NT-2 buffer supplemented with 20mM EDTA pH 8.0, 1mM DTT, and 200U/mL RNase Inhibitor).

Cell lysates were cleared by centrifugation at 20,000G for 10 minutes at 4°C. 100 µL of the supernatant was used for each IP reaction and was added to each tube containing antibody-coated beads. For input, 10 µL was set aside in individual tubes. All tubes were incubated at 4°C with rotation overnight. The following day, the beads were collected using a magnetic rack and were washed in ice-cold NT-2 buffer five times.

For RNA purification, each set of beads was incubated with 150 µL of proteinase K buffer (NT-2 buffer supplemented with 1 % SDS and 1.2mg/ml Proteinase K) at 55°C for 30 minutes while shaking. All input tubes were incubated with 107 µL of NT-2 buffer supplemented with 7.5 µL 20 % SDS, 18 µL of Proteinase K buffer, and 7.5 µL H_2_O at 55°C with shaking for 30 minutes. After digestion, the beads were collected using a magnetic rack and the supernatant was collected and transferred into new tubes. For all samples, including inputs, 250 µL of NT-2 buffer was added before adding 400 µLof acidic phenol:chloroform:isoamyl alcohol (125:24:1) to each tube and vortexing to mix. Samples were centrifuged for 5 minutes at 20,000G to separate phases and the aqueous phase (350 µLl) was removed without disturbing the interphase and transferred into new tubes. To these tubes, 400 µL of chloroform was added and samples were vortexed briefly to mix. Samples were centrifuged again at 20,000G for 5 minutes to separate the phases and the aqueous phase (300 µL) was collected and moved into separate tubes. To each tube, 50 µL of 3M sodium acetate, 15 µL of 7.5M LiCl, 5 µL of 5mg/ml glycogen, and 850 µL of absolute ethanol were added and the samples were incubated for 1 hour at −80°C to precipitate out the RNA.

RNA pellets were collected by spinning the samples at 20,000G for 30 minutes and discarding the supernatant. The RNA pellets were washed in 500 µL ice-cold 80 % ethanol and allowed to air dry at room temp. Dried pellets were resuspended in 20 µL RNase-free H_2_O and stored at -80°C. To test for enrichment of specific RNA species, the isolated RNA was converted into cDNA using the Invitrogen Superscript IV first strand synthesis kit [man: 18091050]. The final product was brought to a total volume of 40 µL. This cDNA was used for comparative CT qPCR using ABI PowerUp SYBR Green Master Mix [cat: A25742] following the included instructions using primers indicated in Supplemental Table 1.

## Results

### *Pantr2* is expressed in both the cytoplasm and nucleus of cortical neural progenitors and their progeny

Knockout of *Pantr2* in a murine model is known to lead to microcephaly (Goff et al., 2015; Sauvageau et al., 2013). This microcephaly is due, in part, to an increase in the thickness and cell number of the deep laminae and a decrease in the cell number and thickness of the upper laminae. The authors also noted a decrease in the production of intermediate progenitor cells, a progenitor pool that is largely destined to generate cells of the upper laminae. This phenotype is summarized in **(Figure 1A)**. To gain a better insight into the changes taking place in the progenitor population in the cortex, we used a cortical neurosphere model to enrich for and culture these progenitors in vitro for further study and analysis.

**Figure 1.**
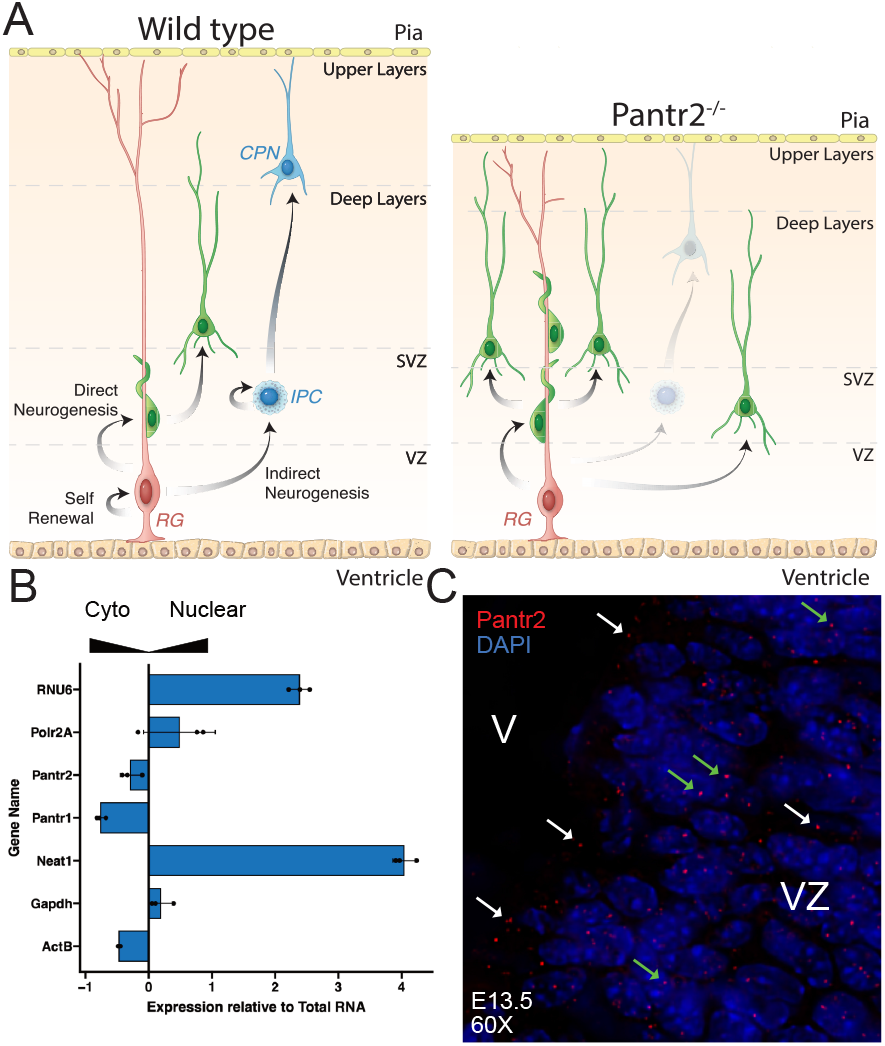
Cartoon depiction of the microcephaly phenotype in the absences of *Pantr2* and the sub cellular localization of *Pantr2*. **(A)** Diagram depicting corticogenesis defects in *Pantr2*^-/-^ mice compared to wildtype. In the wildtype, intermediate progenitor cells (IPCs) are correctly generated from radial glia cells (RGs) in a process known as indirect neurogenesis, and direct neurogenesis continues simultaneously. Conversely, in the *Pantr2*^-/-^ cortex, RGs fail to reliably generate IPCs which contributes to a thickening of the deep layers and a thinning of the upper layers resulting in an overall reduction in cortical thickness.**(B)** qPCR results demonstrating enrichment of known marker genes in either the cytoplasm or the nucleus of wildtype neurospheres.**(C)** Single-molecule fluorescence in situ hybridization (smFISH) against *Pantr2* in E13.5 cortical brain sections taken at 60X magnification and counterstained with DAPI. “SVZ” refers to the subventricular zone, VZ refers to the ventricular zone and RG refers to radial glia cells.

As subcellular location is critical for understanding the mechanism of a lncRNA, we first aimed to determine where *Pantr2* is localized within the cell. To this end, wildtype cortical neurospheres derived from dissociated murine E14.5 dorsal telencephalon were partitioned into nuclear and cytoplasmic fractions, and we used qPCR to quantify the relative enrichment of *Pantr2* transcripts in each fraction compared to whole-cell lysate controls. We observed that *Pantr2* is uniformly expressed in the cytoplasm and nucleus by comparing the enrichment of *Pantr2* to known nuclear-retained RNAs (Neat1 and Rnu6) or RNAs present in both compartments (*Gapdh*) RNAs **(Figure 1B)**. To further validate the subcellular distribution of *Pantr2* RNA, we performed hybridization chain reaction (HCR) mediated single molecule FISH (smFISH). We confirmed the expression of *Pantr2* in the progenitor cells of the ventricular zone (VZ) of wildtype E13.5 cortex as previously described (Goff et al., 2015; Sauvageau et al., 2013) **(Figure 1C)**. Within apical progenitors, we observed *Pantr2* RNA in both the cytoplasm and nucleus **(Figure 1C)**, consistent with the subcellular distribution observed in the fractionation experiment. These results indicate that *Pantr2* transcripts are present in both the cytoplasm and nucleus of cortical progenitors and suggest that *Pantr2* RNA may function within either or both compartments.

### Deletion of *Pantr2* alters the transcriptional landscape of in vitro cortical neurospheres

To identify the gene expression changes that occur in the absence of *Pantr2* we used single-cell RNA sequencing (scRNAseq) of cultured mouse cortical neurospheres. Briefly, primary dissociated cells were collected from the dorsal telencephalon of E14.5 wildtype and *Pantr2*^-/-^ murine embryos and used to derive neurosphere cultures. Dissociated neurosphere cultures (n=2) were used as input for the 10X Genomics Chromium V2 3’ gene expression assay resulting in a dataset containing 25,343 cells. Two clonally-derived populations of neurospheres were used for each genotype, each contributing roughly half of the total number of cells. Data were corrected for batch using a generalized linear model (GLM) (y ~ sex + batch) to account for known sources of technical and biological variation (Haghverdi et al., 2018). Dimensionality reduction for visualization was done using the Monocle3 (Cao et al., 2019; McInnes & Healy, 2018; Qiu et al., 2017; Trapnell et al., 2014) implementation of the uniform manifold approximation and projection (UMAP) algorithm to establish a 2D representation of the first 20 principal components of the batchcorrected dataset **(Figure 2A)**.

**Figure 2.**
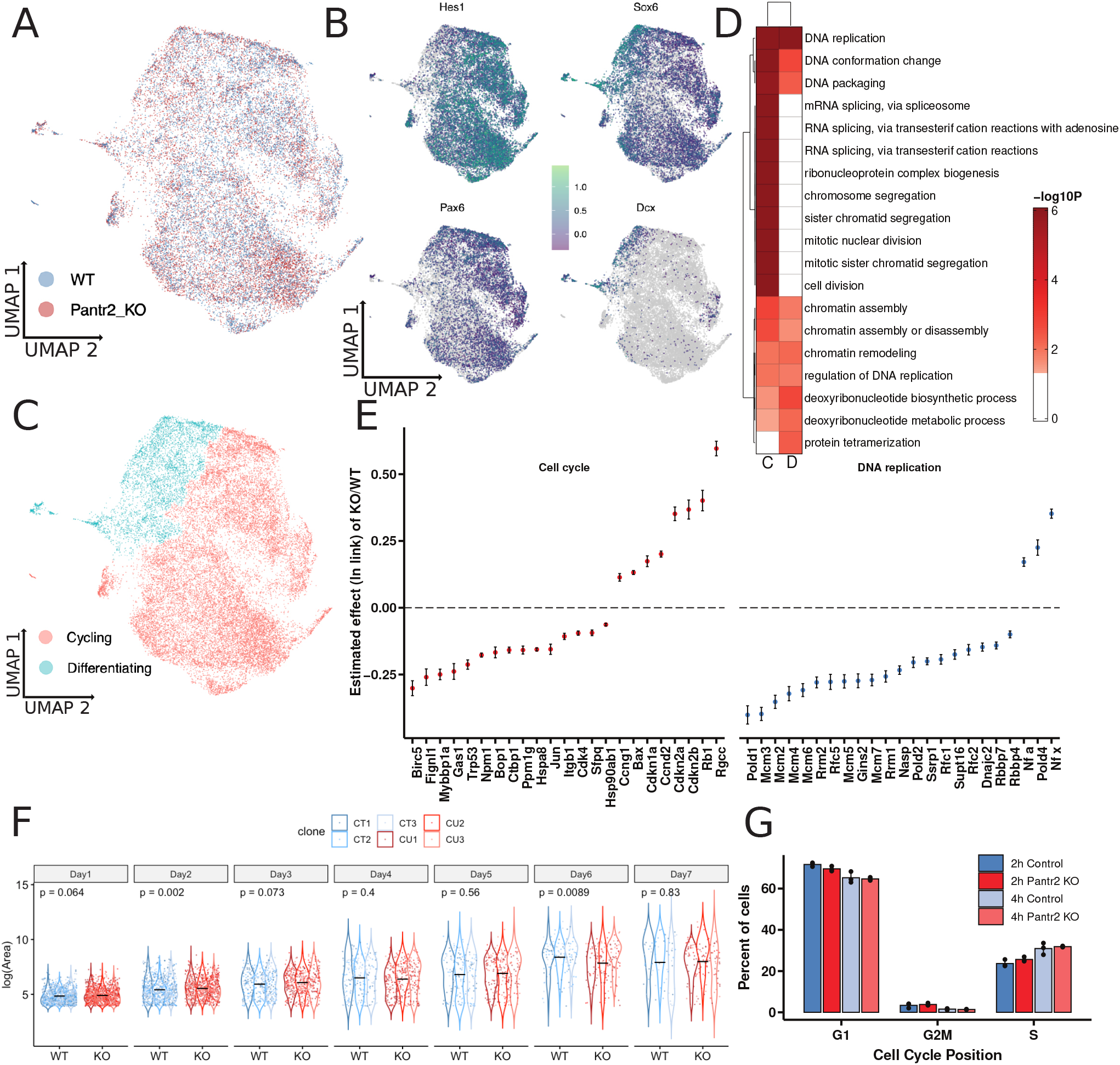
Single cell RNA sequencing for *Pantr2*^-/-^ and Wildtype cortical neurospheres. (**A**) Two-dimensional UMAP representation of wildtype and *Pantr2*^-/-^ neurosphere single-cell RNA sequencing (scRNA-seq) data colored by genotype. (**B**) Log10 expression estimates for Hes1, Sox6, Dcx, and *Pax6* plotted over two-dimensional UMAP representation of wildtype and *Pantr2*^-/-^ neurospheres. (**C**) UMAP representation of wildtype and *Pantr2*^-/-^ colored by cell type. (**D**) Gene ontology enrichment analysis of differentially expressed genes in both the cycling or differentiating progenitors. (**E**) Estimated effect size (ln link) of KO/WT for differentially expressed genes involved in cell cycle regulation and DNA replication. “C” refers to cycling progenitors and “D” refers to differentiating progenitors. (**F**) Calculated areas for wildtype and *Pantr2*^-/-^ neurospheres over seven days. Points are colored by clone ID. (**G**) Percentage of cells found in G1, S, or G2/M of the cell cycle as determined by flow cytometry analysis of a thymidine incorporation assay.

Global differential gene expression analysis between genotypes revealed 887 significantly differentially expressed genes using Monocle3’s implementation of the Wald test on a fitted GLM (y ~ genotype + number of detected genes + sex + batch) **(Supplemental Table 4)**. In the *Pantr2* knockout, we identified upregulated genes involved in the regulation of the cell cycle, including *Ccnd2*, Cdkn2a, and Cdkn2b, suggesting a change in cell cycle dynamics. We also identified the transcription factor Nuclear Factor I X (*Nfix*) and the cell cycle modulator Regulator gene of cell cycle (*Rgcc*) as among the top most significantly-increased genes in *Pantr2*^-/-^ neurospheres by estimated effect size. Expression of *Nfix* is associated with a slowing of the cell cycle, decreased migration of neuroblasts, and differentiation of neuronal progenitors (Heng et al., 2014; Heng et al., 2015; Lu et al., 2020; Zalucki et al., 2019). *Rgcc* is also known to regulate cell cycle through interaction with the mitotic spindle apparatus, usually resulting in an increased G2/M length and an overall longer cell cycle (Saigusa et al., 2007). We also observed significantly differentially expressed genes involved in DNA replication, such as the Mcm family, *Pold1, Gins2, Nsap, Pold2*, and *Rfc2*, that were reduced in *Pantr2*^-/-^ neurospheres. This reduction suggests a disruption of S-phase and further dysregulation of the cell cycle. Notably, the Wnt pathway member *Ndrg2*, known for its role in cell cycle exit (Y. J. Kim et al., 2009), is also increased in the *Pantr2*^-/-^ cells, further implicating a possible abortive exit from the cell cycle, resulting in subsequent neuronal differentiation.

As the generation of intermediate progenitor cells is altered in the *Pantr2*^-/-^ knockout while the abundance of apical progenitors appears unchanged, we hypothesized that the differentially expressed genes list for cells committing to differentiation and those actively cycling would different in response to *Pantr2*. Cells expressing *Hes1* and *Pax6* (Mukhtar and Taylor, 2018) were classified as more progenitor-like than those expressing *Dcx* (Nadarajah and Parnavelas, 2002), a marker of immature neurons **(Figure 2B)**. From this coarse classification, we identified a population of Dcx-expressing, neuronal committed cortical progenitors for further analysis, allowing us to divide the dataset into cycling or differentiating progenitors **(Figure 2C)**. Because the context in which genes are differentially expressed will influence the effect these changes have on cell fate specification, we specifically set out to identify differentially expressed genes (DEGs) between *Pantr2*^-/-^ and wildtype neurospheres in both of the annotated cell states. To this end, we fit a GLM to look at the interaction between genotype and cell state on gene expression (y ~ genotype*assigned cell type + number of detected genes + sex + batch) and performed a Wald test on the combinatorial effect of genotype and assigned cell type. This analysis identified 1255 DEGs (q<5.0×10^−10^) in the actively cycling progenitors and 21 DEGs in the differentiating progenitors **(Supplemental Table 5)**. Gene set enrichment analysis identified significant Gene Ontology (GO) terms associated with DNA replication, cell cycle, and cell division as enriched in the cycling progenitor DEG list, while terms enriched for nucleotide metabolism, RNA degradation, and mRNA processing are enriched within the differentiating progenitor DEG list (Hypergeometric test; p<0.05) **(Figure 2D and Supplemental Table 6)**. Using the estimated effect sizes from the GLM model fit for significant DE genes, we find that most genes associated with DNA replication (GO:0006260) are significantly decreased in the *Pantr2*^-/-^ samples, including *Pold1*/2 (subunits of DNA polymerase) and *Mcm2/3/4/5/7* (components of the minichromosome complex necessary for DNA polymerase elongation), while genes such as *Nfia* and *Nfix* which are associated with a slower cell cycle (Heng et al., 2015; Tchieu et al., 2019), exhibit increased expression in the absence of *Pantr2*. Additionally, genes annotated with the Cell Cycle Regulation GO term (GO:0051726) are decreased overall except for *Cdkn1a, Rb1*, and *Rgcc*, whose expression are associated with slower cell cycle or abortive exit from cell cycle (Giacinti & Giordano, 2006; Karimian et al., 2016; Saigusa et al., 2007) **(Figure 2E)**. In aggregate, these data suggest that loss of *Pantr2* in primary cortical neurospheres leads to a reduction in the cell cycle rate relative to wild-type neurospheres.

Given that we observed changes in the expression of cell cycle-related genes, we next sought to quantify changes in cell cycle dynamics in the absence of *Pantr2* in proliferating cortical neurospheres. To assess this, we used both a neurosphere proliferation assay and a thymidine analog incorporation assay on cultured E14.5 cortical neurospheres. We followed three clonal lines of wildtype neurospheres and three clonal lines of *Pantr2*^-/-^ neurospheres over seven days and measured the average area (in pixels) of individual neurospheres as a proxy for their rate of division **(Figure 2F)**. *Pantr2*^-/-^ neurospheres exhibited a decrease in their average area that reached statistical significance on days 2 and 6. This suggests that cultured *Pantr2*^-/-^ primary neural progenitors have a reduced proliferation rate relative to wildtype. To further test for differential proliferation rates, we pulsed recently-passaged neurospheres with the thymidine analog 5-ethynyl-2’deoxyuridine (EdU) for either 2 or 4 hours and labeled the incorporated EdU with a fluorophore. We performed flow cytometry on each of the clones (n=3) and assigned individual cells to specific phases of the cell cycle based on their relative DNA content **(Figure 2G)**. We tested for differential distribution of cells across cell cycle phases but observed no statistically significant difference between the wildtype neurospheres and *Pantr2*^-/-^ neurospheres maintained in proliferating conditions. These results indicate a change in cell cycle dynamics for *Pantr2*^-/-^ neurospheres when cultured for an extended period (<6 days), however, this difference does not manifest during the shorter time frame examined in our EdU incorporation assay.

### Loss of *Pantr2* alters the chromatin landscape in invitro cortical neurospheres

One potential mechanism through which *Pantr2* may lead to altered gene expression is by altering chromatin accessibility. We performed an assay for transposase-accessible chromatin with next-generation sequencing (ATAC-seq) in cultured E14.5 cortical neurospheres (Buenrostro et al., 2013; Buenrostro et al., 2015). Nuclei were isolated from replicate (n=2) neurospheres from both wildtype and *Pantr2*^-/-^ cultures, and chromatin was enzymatically fragmented using Tn5 transposase. Differential accessibility analysis of the ATAC-seq data between wildtype and *Pantr2*^-/-^ neurospheres identified 847 significantly differentially accessible (DA) peaks using the DiffBind (Ross-Innes et al., 2012) implementation of the Wald test (via DeSeq2) (Love et al., 2014) at a p-value cutoff of p<0.05 **(Supplemental Table 7)**. Of the 847 differentially accessible peaks, 838 were gained in the *Pantr2*^-/-^ indicating overall increase in genomic accessibility in the *Pantr2* KO **(Figure 3A)**. This bias suggests that *Pantr2* is involved in repressing target transcription factors and regulatory elements from binding their DNA target. After annotating peaks based on their distance from the transcription start site (TSS) of neighboring genes, we subset this DA peak list to the 565 peaks within 15Kb-upstream or 3Kb downstream of a known gene TSS **(Supplemental Table 8)**. Using Gene Ontology (GO) enrichment analysis (hypergeometric test; p<0.05) on the genes associated with the differential peaks yielded two enriched GO terms for Biological Processes that reached statistical significance (p<0.05): regulation of cellular response to growth factor stimulus (GO:0090287), and chromosome separation (GO:0051304) **(Supplemental Table 9)**. This suggests a potential functional role for *Pantr2* in mediating epigenetic responses to external growth factor stimuli.

**Figure 3.**
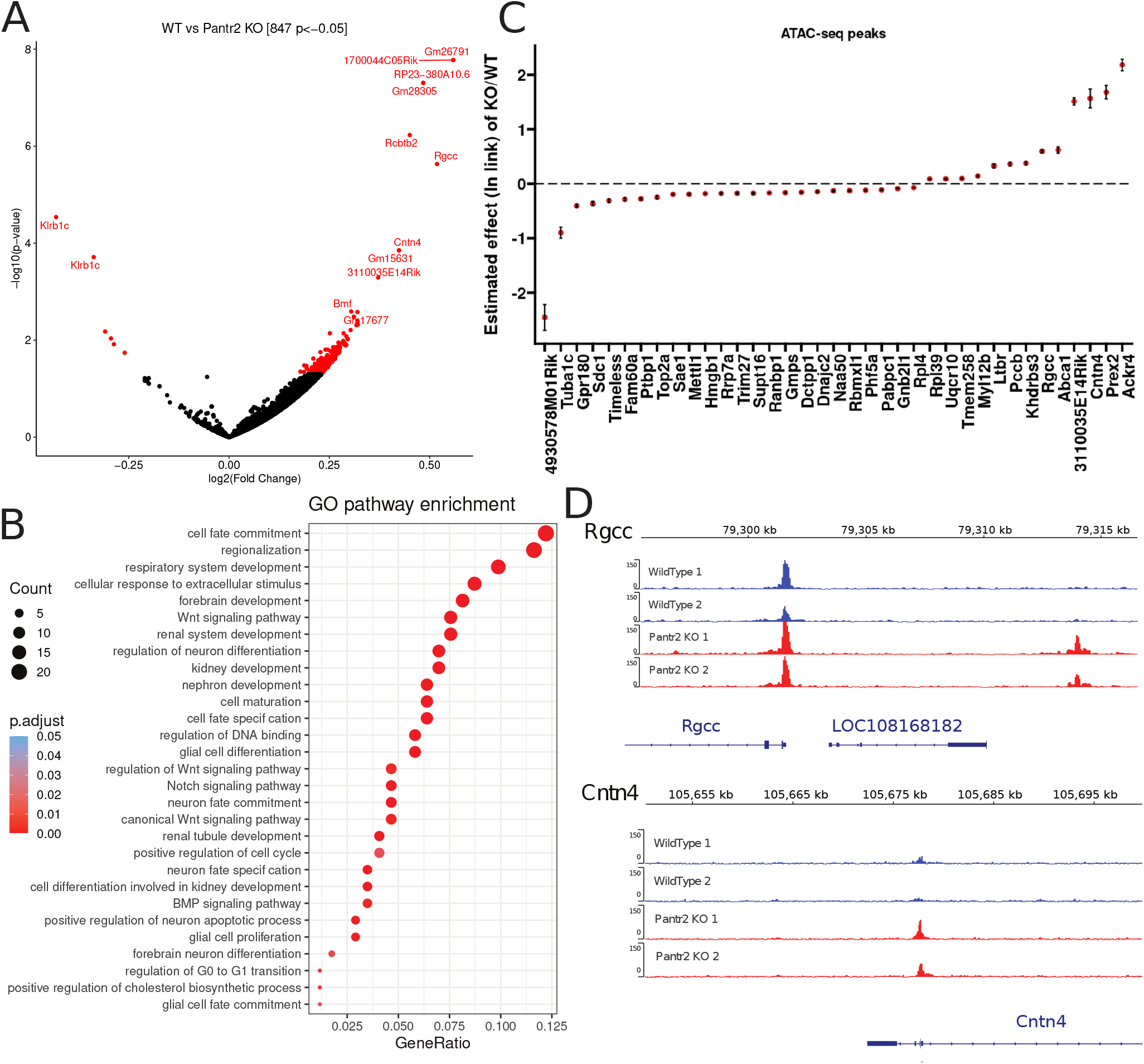
Assay for transposase accessible chromatin paired with next generation sequencing on *Pantr2*^-/-^ and wildtype cortical neurospheres. (**A**) Volcano plot showing all of the differential peaks discovered from the comparison of wildtype and *Pantr2*^-/-^ neurosphere ATAC-seq data. The 847 differential peaks that reached statistical significance are colored in red. (**B**) The estimated effect size of differentially expressed genes from the cortical neuorsphere scRNA-seq dataset that correspond to genes associated with differential peaks from the ATAC-seq analysis. (**C**) Select gene ontologies from GO enrichment analysis for motifs discovered under differential motifs in the ATAC-seq analysis. (**D**) Example bigwig traces for differential peaks associated with *Rgcc* and *Cntn4*, two genes whose expression was also increased in the *Pantr2*^-/-^ neurospheres.

To identify motifs for DNA binding proteins enriched under our differential peaks we used the MEME Suite Analysis for Motif Enrichment (AME) pipeline and identified 217 enriched motifs **(Supplemental Table 10)**. When we intersected the list of DNA binding proteins associated with these motifs with the DNA binding proteins expressed in our scRNA-seq dataset, we identified 112 as downstream candidate genes contributing to the observed differential accessibility. This list includes Kruppel-like factor (KLF) and Specificity Protein (SP) family members as the highest ranked of the significantly enriched motifs. These families of proteins are known to play pivotal roles in stem cell maintenance and differentiation (Presnell et al., 2015). KLF/SP family proteins are also implicated as regulators of Wnt signaling, a process involved in establishing the spatial organization of the brain during development (Patapoutian & Reichardt, 2000). Additionally, we identified several *Pou3f* family member motifs as being enriched in the differential peaks, suggesting that the DNA binding activity of these proteins is affected even if their expression (as determined by scRNA-seq) is not. *Pou3f3* and *Pou3f2* are instrumental for the stereotypical organization of the cerebral cortex (Castro et al., 2006; Dominguez et al., 2013) and the reduction of their affinity to their DNA targets could phenocopy loss of function for these transcription factors(Dominguez et al., 2013; Sugitani et al., 2002). Previous bulk RNA-seq analysis has also suggested that several Pou3f family members (with the specific exception of the neighboring *Pou3f3* gene) are deferentially regulated in the absence of *Pantr2* (Goff et al., 2015). The fact that *Pou3f3* accessibility is impacted in particular is critically important as many lncRNAs have been shown to impact the transcriptional regulation of neighboring genomic loci(Engreitz et al., 2016; Groff et al., 2016; Kopp & Mendell, 2018), while *Pantr2* may instead impact the DNA-binding function of the neighboring *Pou3f3* protein product. Finally, we used GO enrichment analysis to identify the enrichment of motifs associated with GO terms for Biological Processes (hypergeometric test; p<0.05) From this analysis, we found that genes associated with pattern specification (GO:0007389), cell fate commitment (GO:0045165), regionalization (GO:0003002), Wnt signaling pathway (GO:0016055), Notch signaling pathway (GO:0007219) and forebrain development (GO:0030900) GO terms were enriched for from our list of enriched motifs **(Figure 3B and Supplemental Table 11)**.

Using the scRNA-seq data collected from wildtype and *Pantr2* KO neurospheres, we next asked whether the expression of any genes neighboring differential peaks also exhibited significant differential expression. 37 genes were found to have significant differential expression (Wald test; q<5.0×10^−10^), potentially as a result of differential usage of regulatory elements associated with these genomic loci **(Figure 3C)**. Of these DEGs, two are relevant to the *Pantr2* KO phenotype **(Figure 3D)**; Cntn4 is associated with cell-cell adhesion and aspects of neuronal maturation such as axonal outgrowth and dendrite formation (Fernandez et al., 2004), and *Rgcc* is a regulator of cell cycle, a process known to be linked with differentiation potential in the developing cortex. These findings demonstrate differential accessibility after KO of *Pantr2*, suggesting that the observed phenotypes may result from epigenetic dysregulation as a consequence of *Pantr2* loss, and provide targets such as the KLF/SP family and the *Pou3f* family of transcription factors.

### *Pantr2* is a trans-acting lncRNA that promotes proliferation in differentiating neural precursors

We have demonstrated that the *Pantr2* locus is required for proper corticogenesis and laminar specification, however, an important question to address when assessing the roles for a lncRNA gene is its functional mechanisms. To directly assess whether the *Pantr2* transcript itself is biologically active, we cloned the *Pantr2* cDNA into a previously described lncRNA expression vector (Hacisuleyman et al., 2016) and ectopically expressed the lncRNA gene in mouse Neuro2a (N2a) cells. We first demonstrated that *Pantr2* could be efficiently over-expressed in N2a cells and that this increase in expression does not impact the expression of the neighboring *Pou3f3* gene **(Figure 4A)**. In a naive, undifferentiated state, *Pantr2* RNA expression alone was insufficient to significantly alter the expression of both *Nfix* and *Rgcc*, both of which were significantly upregulated in the *Pantr2*^-/-^ neurospheres **(Figures 4B and 4C)**. We, therefore, hypothe-sized that any observed phenotype associated with changes in *Pantr2* expression may be restricted to a window surrounding the commitment to neurogenic differentiation into neural precursor cells. To test whether ectopically-expressed *Pantr2* affects the expression of previously identified target genes under differentiating conditions, N2a cells were transfected with *Pantr2* expression constructs and stimulated to differentiate via serum withdrawal. After stimulation of differentiation for 48 hours, *Pantr2* over-expressing N2a cells showed a significant decrease in *Rgcc* expression relative to empty-vector controls **(Figure 4C)**; a complementary result to that observed in the *Pantr2*^-/-^ neurosphere scRNA-seq dataset. We also examined the expression of the NFI family of transcription factors, as they appeared to be sensitive to *Pantr2* expression, and found that only under differentiating conditions were *Nfia* and *Nfix* significantly suppressed in response to *Pantr2* over-expression **(Figure 4C)**. To assess the impact of *Pantr2* over-expression on neurogenic differentiation, we next evaluated the expression of select genes involved in neurogenesis and found that *Ngn2, Notch1*, and *Fbxw7* were significantly reduced in the *Pantr2* over-expression under differentiating conditions **(Figure 4D)**. The observed reduction in *Ngn2* is particularly interesting, as this gene is known to be induced by N2a cells during differentiation (Muñoz et al., 2003). These results suggest that expression of *Pantr2* under induced differentiating conditions can act as an inhibitor of differentiation in trans, a complementary phenotype to that observed in vivo in the Pantr2^-/-^ mouse. Interestingly, this effect is not observed in N2a cells maintained under proliferating conditions, suggesting that the functional consequences of changes in *Pantr2* gene expression may be less pronounced in progenitor cells, and stronger in cells undergoing commitment to a neuronal fate. This suggests that the specific functional role(s) for *Pantr2* are context-specific. The ectopic expression of *Pantr2* RNA in differentiating N2a cells elicits a reproducible effect on key genes associated with neuronal differentiation, demonstrating a trans-acting biological function for the *Pantr2* RNA in regulating gene expression during neuronal differentiation.

**Figure 4.**
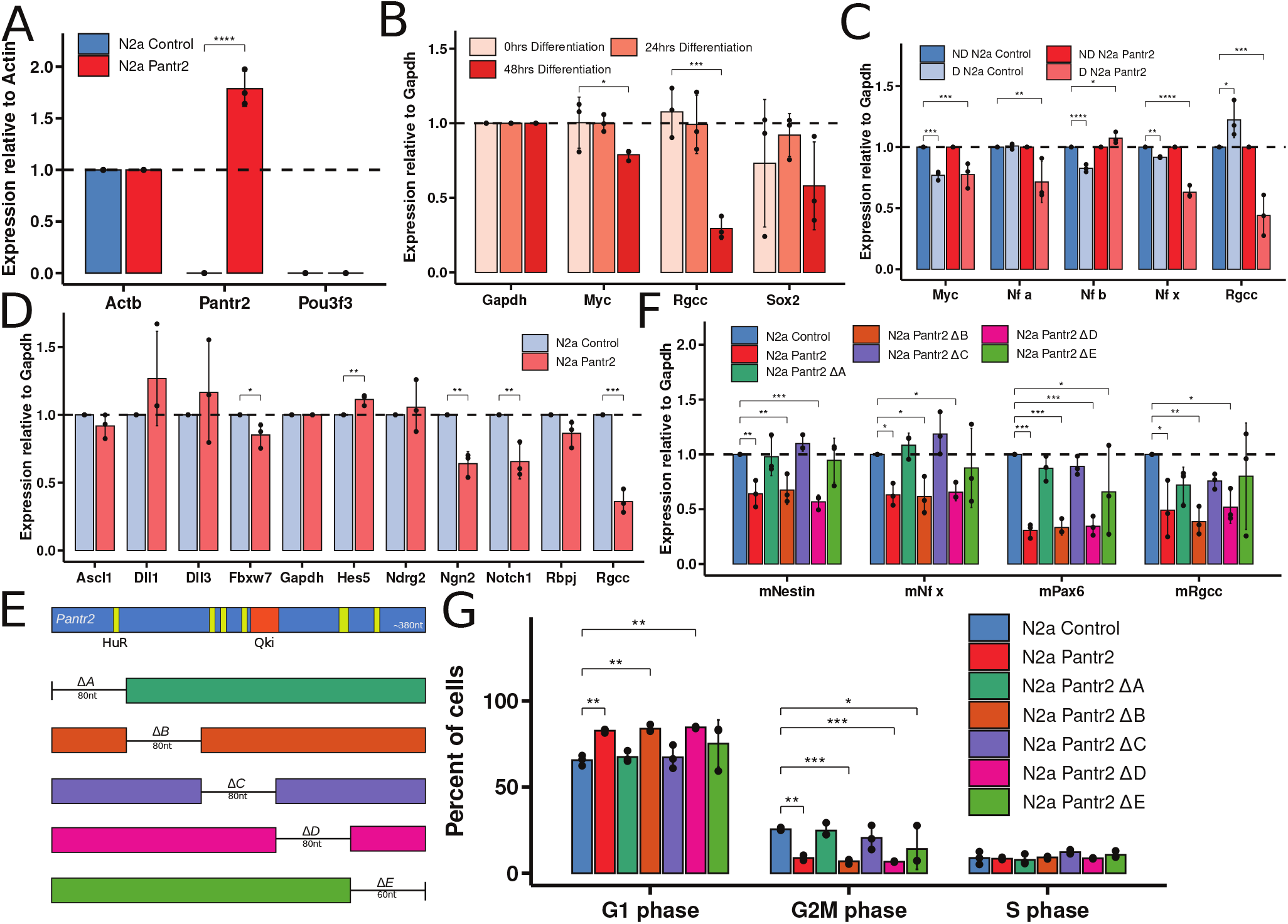
Gene expression analysis of the overexpression of *Pantr2* in N2a cells and the effects on cell cycle progression. (**A**) qPCR results showing the expression of *Pantr2* and *Pou3f3* relative to Actb in N2a cells. (**B**) qPCR results showing the expression of *Myc, Rgcc*, and Sox2 relative to *Gapdh* in N2a cells after 0, 24, and 48 hours of differentiation while ectopically expressing *Pantr2*. (**C**) Comparison of the relative expression of *Myc, Nfia, Nfib, Nfix*, and *Rgcc* in undifferentiated N2a cells and N2a cells differentiated for 48 hours using *Gapdh* as an internal control and N2a empty transfection as the treatment control. “ND” refers to non-differentiation conditions and “D” refers to differentiation conditions. (**D**) qPCR results show the relative expression of several neurogenic genes compared to *Gapdh* in differentiating N2a cells expressing *Pantr2* and an empty control using *Gapdh* as an internal control and N2a empty transfection as the treatment control. (**E**) Deletion mapping scheme for *Pantr2* truncations and annotation of HuR and Qki sites. (**F**) qPCR results for the relative expression of *Nestin, Nfix, Pax6* and *Rgcc* compared to *Gapdh* in differentiating N2a cells expressing *Pantr2, Pantr2*Δ A, *Pantr2*Δ B, *Pantr2*Δ C, *Pantr2*Δ D, *Pantr2*Δ E or empty control using *Gapdh* as an internal control and N2a empty transfection as the treatment control. (**G**) Percentage of cells found in G1, S or G2M of the cell cycle as determined by flow cytometry analysis of a thymidine incorporation assay in differentiating N2a cells expressing *Pantr2, Pantr2*Δ A, *Pantr2*Δ B, *Pantr2*Δ C, *Pantr2*Δ D, *Pantr2*Δ E or empty control. All qPCR data was normalized by setting the internal control and the treatment control (both specified per figure panel) to 1.

Having demonstrated that *Pantr2* over-expression affects the expression of key genes involved in neurogenesis in differentiating N2a cells, we next asked whether we could identify functional domains of the *Pantr2* RNA that might be contributing to this trans-acting effect. To this end, we used deletion mapping to generate a panel of *Pantr2* deletion mutants for analysis. Using site-directed mutagenesis, we created 5 unique deletion constructs (*Pantr2* Δ A - Δ E) in which 80nt regions were removed in each except for *Pantr2* Δ E, in which the last 60nt of *Pantr2* were removed **(Figure 4E)**. For this assay, we define *Pantr2* activity as the ability to inhibit the expression of *Pantr2* responsive genes in differentiation conditions as measured by qPCR. Each deletion mutant was independently expressed in N2a cells under differentiating conditions to determine which regions were critical for mediating the observed changes in *Rgcc* and *Nfix* expression after *Pantr2* over-expression. We observed that *Pantr2* Δ A, Pantr2 Δ C, and *Pantr2* Δ E mutants lost *Pantr2* activity **(Figure 4F)**, suggesting that these regions harbor functional elements of the *Pantr2* RNA.

To identify potential functional sequence motifs within these deleted regions, we annotated putative RNA binding protein (RBP) motifs contained within the *Pantr2* Δ A, *Pantr2* ΔC, and *Pantr2* Δ E regions annotated using the RNA binding protein database RBPDB (Cook et al., 2011). We identified multiple binding sites for HuR, an RBP encoded by the Elavl1 gene, within the *Pantr2* Δ A, *Pantr2* Δ C, *Pantr2* Δ D, and *Pantr2* Δ E regions of *Pantr2* **(Figure 4E)**. Previous studies have described an interaction between an isoform of *Pantr2* and HuR, further implicating this RBP as a potential mediator of *Pantr2*-dependent epigenetic regulation of gene expression (Carelli et al., 2019). We also identified an annotated binding site for the RBP QKI in the C fragment of *Pantr2* from Starbase (J.-H. Li et al., 2014) **(Figure 4E)**. As QkI is selectively expressed in apical progenitors during development (Hardy et al., 1996; Hayakawa-Yano et al., 2017), this RBP is also a likely interacting partner for *Pantr2* in this context.

To examine potential cell cycle effects from *Pantr2* expression modulation in N2a cells, we used a thymidine analog (EdU) incorporation assay to probe the proportion of *Pantr2*-expressing N2a cells present in G1, S, and G2/M phases of the cell cycle using flow cytometry. We ectopically expressed full-length *Pantr2*, or the deletion constructs *Pantr2* Δ A-Δ E and an empty control vector in N2a cells. After 48 hours, the cells were changed into differentiation media and allowed to differentiate for 48 hours. After this interval, we performed an EdU incorporation assay to determine the proportion of cells in each cell cycle state **(Figure 4G)**. Following over-expression of *Pantr2*, N2a cells are significantly more likely to reside in G1 phase and less likely to be in G2/M when compared to the control, while the number of cells in S phase is unaffected. This indicates a potential block at the transition from G2/M phase to G1 phase in the control cells during differentiation that the *Pantr2*-over-expressing cells are able to escape. This is consistent with the expected phenotype associated with the observed decreased expression of *Rgcc* and *Nfix* in the *Pantr2*-over-expressing N2a cells during differentiation. When we examine the *Pantr2*Δ A-Δ E mutants, we find that *Pantr2*Δ A, *Pantr2*Δ C, and *Pantr2*Δ E again match the control N2a cells in the proportion of cells present in each of the cell cycle stages, while *Pantr2*Δ B and *Pantr2*Δ D appear to match the cell proportions present in the cell cycle stages of *Pantr2* expressing N2a cells, reinforcing the idea that the *Pantr2*Δ A, *Pantr2*Δ C, and *Pantr2*Δ E mutants have lost *Pantr2* activity, which is retained in the *Pantr2*Δ B and *Pantr2*Δ D mutants. Together, these results demonstrate the ability for the *Pantr2* RNA to modulate the expression of key genes associated with neuronal commitment and cell cycle in trans through ectopic expression. We also validate changes in cell cycle dynamics through over-expression of *Pantr2* RNA and identify possible RNA binding protein motifs that could be required for *Pantr2* activity.

### In vivo loss of function of *Pantr2* affects differentiating neuron fate commitment and the choice of direct vs indirect neurogenesis

Our previous findings suggest that *Pantr2* is a trans-acting lncRNA that largely disrupts DNA binding proteins from their target loci and impacts cell cycle and differentiation potential in vitro. We next asked whether these effects are observed during in vivo corticogenesis in the absence of *Pantr2* and contribute to the previously described mislamination and fate specification defects. To test this, we performed single nuclei RNASeq (snRNA-seq) on murine E15.5 wildtype and *Pantr2*^-/-^ dorsal telencephalon (n=4). After filtering low-quality droplets and doublets (see methods) we were left with 65,319 nuclei for analysis. A UMAP embedding was generated **(Figure 5A)** and Leiden community detection was used to assign cells to six clusters indicated as Groups 1-6 **(Figure 5A)**. Expression of classical cortical developmental markers such as *Hes5, Gli3, Eomes, Satb2, Tbr1, Foxp2, Tle4, and Bcl11b* was used to determine the overall direction of differentiation within the contiguous manifold from apical progenitors in Group 3 to maturing cortical excitatory projection neurons Groups 1, 2, 5 and 6 **(Figure 5B)**. Using these marker gene expression profiles, we further refined the cluster annotations, with Group 3 identified as *Hes5*^+^ apical progenitors and Group 4 as *Eomes*^+^ intermediate progenitor cells (IPCs). Groups 1, 2, 5, and 6 correspond to maturing neurons developing towards deep layer excitatory projection neuron identities, as indicated by high expression of *Tle4*, Foxp2, and Necab1 in Group 2, or more superficial layer excitatory projection neurons as indicated by the expression of *Satb2, Cux1, Lhx2*, and *Pou3f2* in Group 1.

**Figure 5.**
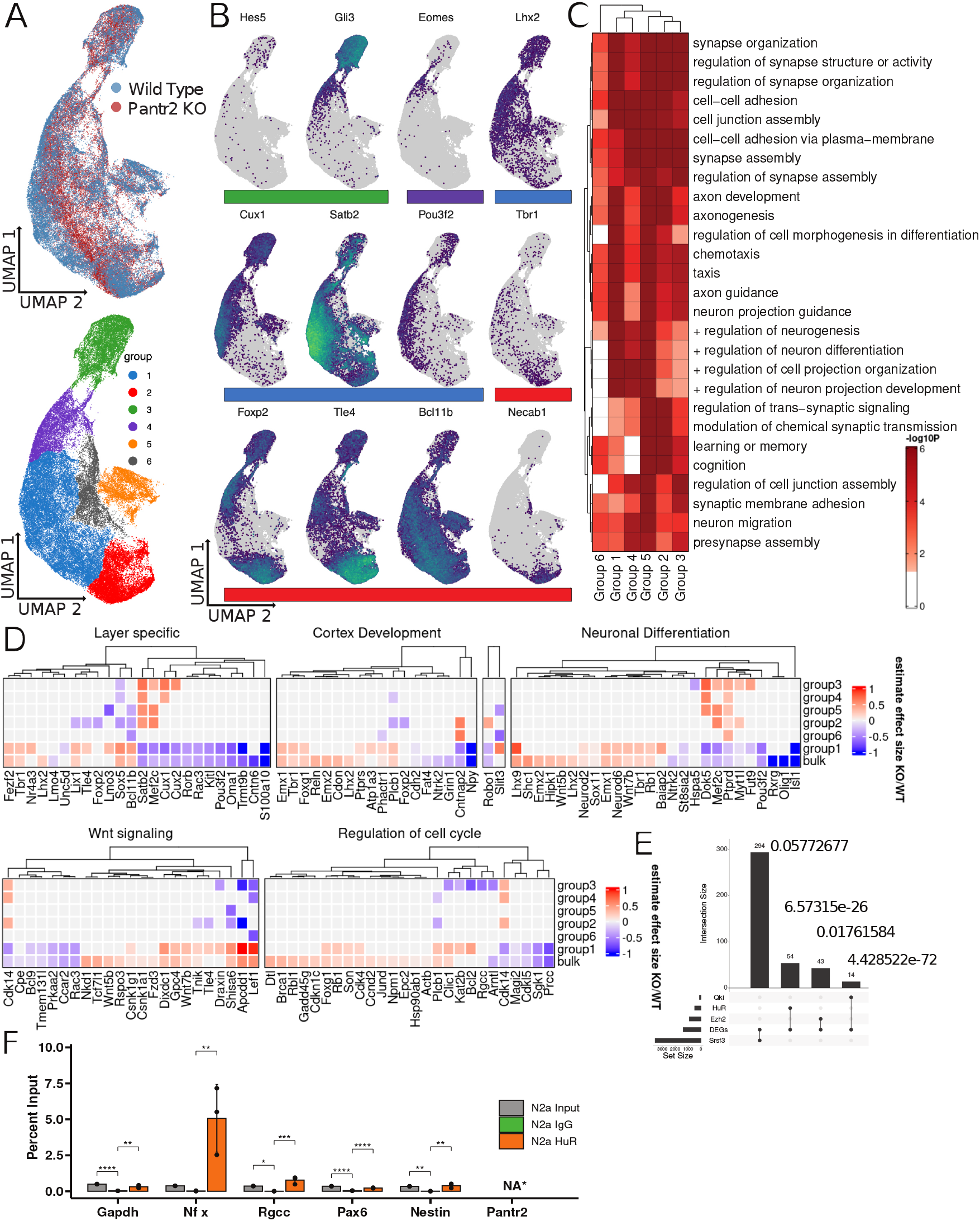
Single nuclei RNA sequencing of *Pantr2*^-/-^ and wildtype E15.5 dorsal telencephalon. (**A**) Two-dimensional UMAP representation of snRNA-seq data from E15.5 mouse dorsal telencephalon colored by genotype (wildtype and *Pantr2*^-/-^) and Leiden community detection identified groups. (**B**) Expression of layer-specific marker genes ordered by expression in groups identified in panel A. (**C**) Gene ontology enrichment analysis on the differentially expressed genes by group colored by -log_10_ p-value. (**D**) Heatmap of estimated effect size from the differentially expressed genes by genotype in bulk and in each group. Genes were chosen by gene ontology terms and a list of layer-specific genes from Molyneux et al 2009. (**E**) Upset plot showing the intersection of the differentially expressed genes found in the snRNA-seq dataset and publicly available CLIP-seq data for the RNA binding proteins QKI, HuR, EZH2, and SRSF3. Each intersection is annotated with a p-value from the hypergeometric test performed to see if their is enrichment of RBP targets in the differential genes list. (**F**) RNA immunoprecipitation (RIP) of HuR coupled with qPCR to detect binding of *Pantr2, Nfix, Rgcc, Gapdh, Pax6* asnd *Nestin* to the RBP HuR in N2a cells. Samples are compared both to an input and an IgG negative control.

To identify gene expression differences that may be specific to different states of neuronal maturation, we performed clusterspecific differential gene expression analysis using a Wald test on a fitted GLM (y ~ genotype*group + number of detected genes) to identify the significant combinatorial effects of genotype and cluster on gene expression. Using this method, we were able to identify 1,420 differentially expressed genes with respect to cluster **(Supplemental Table 12)**. Unexpectedly, there are only 25 genes (2.3 %) that are jointly differentially expressed with respect to genotype in our neurosphere dataset and with respect to genotype in Group 3 highlighting the differences between our in vivo and in vitro models **(Supplemental Figure 3)**. Group 3 is especially relevant as it represents apical progenitors, a cell type that is most similar to those reflected by cortical neurosphere generation. GO enrichment analysis on the cluster-specific DEGs revealed terms associated with cell adhesion molecules (GO:0007155), positive regulation of neurogenesis (GO:0050769), neuron differentiation (GO:0045664), and neuron migration (GO:0001764); each relevant in regulating fate specification of apical progenitors **(Figure 5C and Supplemental Table 13)**. Cortical layer-specific genes, as defined by Molynaeux et al 2007, are also significantly differentially expressed between *Pantr2*^-/-^ and wild-type nuclei as anticipated given the previously-described lamination defects **(Figure 5D)** (Sauvageau et al., 2013). For example, marker genes associated with upper-layer callosal projection neurons such as *Satb2, Cux1, Mef2c, Rorb*, and *Pou3f2* are decreased in the *Pantr2* KO while genes associated with deep layer projection neuron subtypes such as *Tbr1*, Lhx2, *Tle4*, and *Bcl11b* are increased in both Group 1 and the pseudo-bulk **(Figure 5D)**. Interestingly, several markers for upper-layer cortical projection neurons including *Satb2, Mef2c, Cux1*, and *Cux2* are increased in the *Pantr2*^-/-^ in Group 3 apical progenitors, despite this population being underrepresented in the maturing neuron states **(Figure 5D)**.

Many genes involved in Wnt signaling are dysregulated in the *Pantr2* knockout dorsal telencephalon **(Figure 5D)**. Specifically, Wnt antagonists such as *Tle4, Nkd1, Shisa6, Apcdd1, and Draxin* are increased and agonists such as,*Lef1, Rspo3, Tnik*, and *Dixdc1* are also upregulated in the *Pantr2*^-/-^ telencephalon at this time point. Among the downregulated genes we also find the Wnt agonists *Bcl9, Rac3*, and *Prkaa2*, and antagonists such as *Cdk14* and *Cpe*. Despite the mixed representation of positive and negative regulators and effectors of Wnt, the significant (p.adj = 5.87e-05, hypergeometric test) over-representation of Wnt-associated genes suggests that Wnt signaling, a key process in cortical neuron differentiation and maturation (A. S. Kim et al., 2001; Munji et al., 2011), is likely dysregulated.

Likewise, several genes involved in cell cycle regulation are also differentially expressed **(Figure 5D)**, again indicating defects in cell cycle dynamics in the *Pantr2* knockout. For example, the DNA damage checkpoint regulator *Dtl* is upregulated in the *Pantr2*^-/-^ which is associated both with the regulation of exit from S-phase and passage through G2/M. Increased expression of *Dtl* is seen in some cancers and is associated with an increase in proliferation. Also, *Dtl* levels decrease during induced differentiation in HepG2 and NT2 cell lines indicating involvement in the differentiation process (Pan et al., 2006). Increased *Bcl2* expression, which is associated with a block at the G1/S phase transition (Zinkel et al., 2006), is observed in immature neurons expressing upper-layer markers (Group 1) but decreased in the apical progenitors (Group 3), potentially indicating a change in cell cycle dynamics in the penultimate cycle and neurogenic fate commitment. Additionally, *Rb1* has increased expression relative to the control, indicating a potential decrease in cell cycle rate (Giacinti & Giordano, 2006). As RB1 is phosphorylated, thus inactivated, by *Ccnd2* (Wang et al., 2016) it is surprising that *Ccnd2* expression is also increased in the knockout, but is consistent with our findings from the scRNA-seq of *Pantr2*^-/-^ neurospheres **(Figure 2E)**.

We annotated putative QKI and HuR binding sites present on the *Pantr2* transcript and demonstrated that these sites were present on regions of *Pantr2* that are required for its function **(Figure 4E)**, suggesting that interactions with these proteins may mediate some of the observed downstream effects of *Pantr2* activity. With a list of significantly differentially expressed genes impacted by the loss of *Pantr2* in an in vivo developmental context in hand, we next assessed if known QKI- or HuR-binding genes are enriched in this set. For this analysis, we chose EZH2 and SFRS3 as negative controls, as EZH2 is found bound to many transcripts promiscuously and SFRS3 is a housekeeping RBP that does not appear to be preferential to tissue or cell type. The intersection of all DEGs with either QKI, HuR, EZH2, or SFRS3 target genes are provided in **(Figure 5E)**. A hypergeometric test was used to test for enrichment of these RBP target genes in our DEG list. We identified a significant enrichment of QKI and HuR target genes in our differentially expressed genes (p = 4.428e^−72^ and p = 6.573e^−26^ respectively), indicating the potential involvement of these RBPs in mediating the downstream effects of *Pantr2*^-/-^. We found no significant enrichment for SFRS3 or EZH2 binding sites (p = 0.057, p = 0.017) indicating specificity for QKI and HuR targets over SFRS3 and EZH2 as hypothesized.

To test if *Pantr2* is bound to HuR in vitro, we next utilized a native RNA immunoprecipitation (RIP) assay to precipitate HuR protein from N2a lysate and probe for associated RNA molecules using qPCR **(Figure 5F)**. We found that, while *Nfix* is indeed enriched in RNAs isolated from the HuR RIP as previously reported (Rothamel et al., 2021), *Pantr2* was not enriched in this population when compared to other genes not reported to be associated with HuR, indicating that it is unlikely that *Pantr2* is directly associated with HuR in this context.

Finally, to further explore the effect on *Pantr2* knockout on cell cycle in both our scRNA-seq neurosphere dataset and the snRNA-seq dorsal telencephalon dataset, we annotated cell cycle position using the R package tricycle (Zheng et al., 2022). Using this method, we looked for deviations in cell cycle states between the two genotype conditions. We identified a significant (Wilcoxon Rank Sum Test) shift in cell cycle state between the *Pantr2*^-/-^ and wildtype cortical neurospheres, consistent with altered cell cycle state occupancy **(Supplemental Figure 2A and 2B)**. When similarly applied to the snRNA-seq dorsal telencephalon dataset, tricycle analysis of each of the annotated clusters **(Figure 5A)** identified a significant difference in cell cycle state occupancy in Group 3 (apical progenitors), consistent with an increased number of *Pantr2*^-/-^ cells present in G2/M, as predicted based on the changes in *Rgcc* expression **(Supplemental Figure 2A and 2)**. In aggregate, these results provide evidence of cell cycle defects present in the apical progenitors of the *Pantr2*^-/-^ dorsal telencephalon, consistent with our in vitro observations.

### Differential fate specification in *Pantr2* KO mouse corticogenesis

We next asked whether we could identify differences in the differentiation fate potential of neuronal progenitors in the *Pantr2* KO. We theorized that any defects in fate specification arising from the loss of *Pantr2* in the E15.5 snRNA-seq data would result in differential abundance of cells by genotype across the observed differentation manifold **(Figure 5A)**. To identify subgraph regions that are biased for the abundance of cells by genotype, we used miloR (Dann et al., 2022) to group cells into neighborhoods using the underlying KNN graph embedding and examine the differential contribution of each genotype to the composition of groups of neighborhoods. In this manner, we generated unique neighborhood groups that are learned independently from the previously annotated clusters, and identified neighborhood groups with significant enrichment for cells of either genotype.

Consistent with our previous observations that actively differentiating cells appear to have a more pronounced transcriptional phenotype in the *Pantr2*-/- mice, we observed minimal differential abundance with respect to genotype in apical progenitors, and a significant differential abundance of *Pantr2*^-/-^ cells in neighborhoods associated with neuronal commitment and maturation **(Figure 6A)**. Specifically, we identify that neighborhood groups 9 and 8 are significantly enriched for *Pantr2*^-/-^ cells while neighborhood groups 6 and 12 are significantly depleted **(Figure 6B)**. Further, we identified marker genes for these neighborhoods and used these markers to identify changes in fate specification programs unique to each genotype. For example, *Nfib, Nfia*, and *Sox5* are more highly expressed in the *Pantr2*^-/-^ -enriched neighborhood groups, while *Lsamp, Robo1*, and *Robo2* expression mark the neighborhood groups containing significantly more wildtype cells **(Figure 6C)**. When we aggregate marker genes for genotype-biased neighborhoods, we observe that the markers for neighborhoods over-represented by *Pantr2*^-/-^ cells are enriched for GO terms corresponding to cell cycle **(Figure 6D)**. In contrast, the markers for neighborhoods depleted for *Pantr2*^-/-^ cells are enriched for GO terms related to neuronal maturation. These results suggest that there is a change in the cell fate trajectories of differentiating *Pantr2*^-/-^ cells, resulting in part from disruption to the normal cell cycle dynamics, and contributing to the laminar identities of the developing cortical projection neurons.

**Figure 6.**
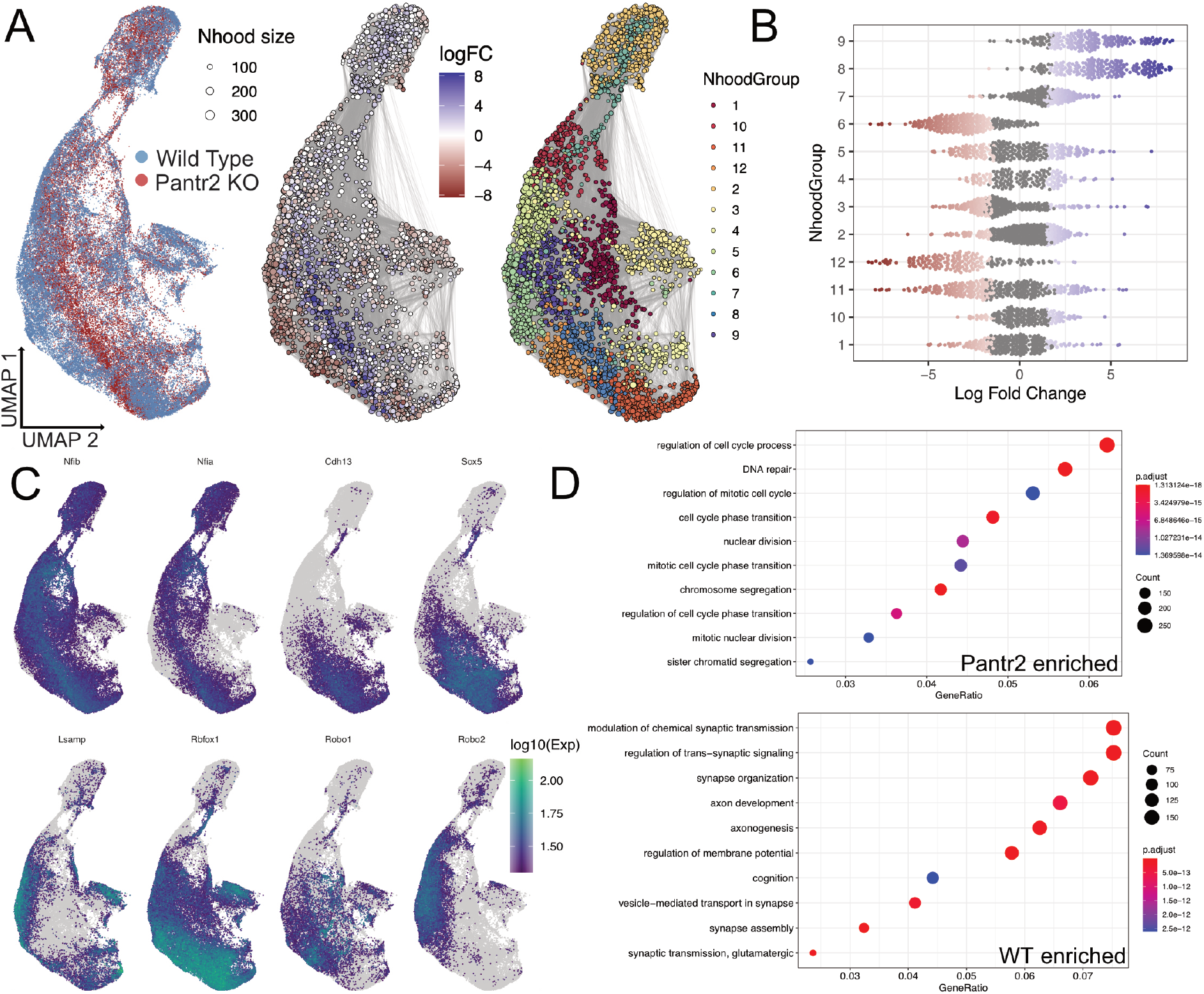
Differential abundance analysis for single nuclei RNA sequencing of *Pantr2*^-/-^ and wildtype dorsal telencephalon. (**A**) Two-dimensional UMAP representation of snRNA-seq data from E15.5 mouse telencephalon colored by genotype (wildtype and *Pantr2*^-/-^), miloR identified differential abundance by genotype and identified neighborhood groups identified by miloR. (**B**) Beehive plot for the proportion of neighborhoods by genotype within each neighborhood group. (**C**) Sample marker genes for the neighborhood groups with differential abundance of neighborhoods by genotype. (**D**) Gene ontology enrichment analysis of marker genes for both the *Pantr2*^-/-^ enriched neighborhood groups and wildtype (WT) enriched neighborhood groups.

## Discussion

Previous work identified *Pantr2* as a lncRNA involved in the stereotypical organization of the cortex during development by demonstrating that loss of this lncRNA is sufficient to cause cortical lamination defects and microcephaly. This study extends this by identifying specific pathways that are perturbed in the absence of this key lncRNA gene. We demonstrate that *Pantr2* acts in trans and exerts its effects principally under differentiation conditions in vitro. We also demonstrate cell cycle alterations attributed to loss or gain of function of *Pantr2* that are consistent across models. We identify two *Pantr2*-responsive genes, *Nfix* and *Rgcc*, as likely contributors to the observed *Pantr2* knockout phenotype. Increased expression of *Nfix* has been shown to promote exit from the cell cycle and is consistent with the observed precocious differentiation in the *Pantr2*^-/-^ cortex(Clark et al., 2019; Heng et al., 2015; Zalucki et al., 2019),. While the function of *Rgcc* is not fully understood in the context of neural development, it is known to play a role in regulating the length of M-phase. Specifically, increased expression of RGCC was noted to increase the length of M-phase (Saigusa et al., 2007). Recent studies have tied cell cycle length increase, especially in M-phase, to precocious apical progenitor differentiation in several models of neuronal differentiation (Mitchell-Dick et al., 2019; Pilaz et al., 2016). In the context of *Pantr2* loss of function (LOF), precocious differentiation is consistent with the observed expansion of deep layer neurons at the expense of upper layers, as deep layers are generated first during corticogenesis. Together, our data suggest that *Pantr2* promotes proliferation and expansion of radial glia while suppressing differentiation and that loss of this lncRNA results in a slower cell cycle and an abortive exit of cell cycle that likely results in precocious differentiation into neurons.

In neurospheres, we found that loss of *Pantr2* results in widespread transcriptional changes and changes in chromatin accessibility. In the absence of *Pantr2*, we see enrichment of inaccessible regions marked with *Pou3f3, Pou3f2*, and KLF/SP family motifs indicating potential involvement of the Wnt signaling pathway and Pou3f family members. Differential gene expression analysis in neurospheres shows that *Pou3f3* appears to be unchanged between the *Pantr2*^-/-^ and wildtype suggesting that the changes we see in motif utilization are not due to changes in mRNA expression and could be due to changes in protein level or function. Moreover, the gene expression changes that we identified in our scRNA-seq neurosphere dataset suggest that the largest difference between *Pantr2*^-/-^ and wildtype neurospheres are the regulation of processes relating to cell cycle. This is very intriguing as several reports have linked cell cycle dynamics to differentiation potential in the developing cortex (Mitchell-Dick et al., 2019; Pilaz et al., 2016; Yeh et al., 2014). We see evidence of slower cell cycle progression in our neurosphere assay but not in our thymidine incorporation assay. This finding led us to conclude that either 2-4 hours is not long enough to see an effect in neurospheres or that the neurosphere assay confounds proliferation and differentiation and is thus not an ideal model for isolating cell cycle effects in a time course.

Using an over-expression system in N2a cells, we demonstrated that expression of *Pantr2* exerts its effect specifically in differentiation conditions, making the N2a cells more resistant to differentiation. This is evident by the reduced expression of *Ngn2* in differentiating N2a cells expressing *Pantr2*. We also see evidence of a faster cell cycle in the presence of *Pantr2* in the N2a cells, seeing fewer cells in G2/M when compared to G1 with no change in the number of cells captured in S-phase, as measured by thymidine incorporation assay. This indicates a pileup of cells at G2/M in differentiating N2a cells that are relieved in response to *Pantr2* expression. We also demonstrate that gain of *Pantr2* in N2a cells leads to reduced expression of *Nfix* and *Rgcc*. Consistent with this, loss of *Pantr2* in neurospheres increased both genes indicating that these two genes are repressed by *Pantr2* expression. Lastly, we used deletion mapping of *Pantr2* to determine functional domains of *Pantr2* and found three independent regions (60-80nt) that are required for *Pantr2* activity. This result further demonstrates a functional element of the lncRNA itself as being responsible for the activity of *Pantr2*.

We utilized snRNA-seq to look for gene expression changes in the embryonic dorsal telencephalon at E15.5 to validate our findings in vivo. We identify differentially expressed genes that are shared between the neurosphere dataset and this embryonic dataset, highlighting *Nfix* and *Tcf4* for their roles in corticogenesis. We were also able to determine that most of the changes in gene expression were found in immature neurons, especially those expressing *Pou3f2* and *Satb2*, both known markers for upper-layer neurons. Interestingly, *Pantr2* knockout dorsal telencephalon cells have reduced expression of upper layer markers and increased expression of deep layer markers, providing some evidence that the predominant *Pantr2*^-/-^ phenotype is one of precocious differentiation rather than neuronal migration. We provide further support to this model by identifying cell cycle dynamics as being a major source of deviation during early neurogenesis.

Lastly, we explored a potential interaction between *Pantr2* and the RNA binding protein HuR. Previous work has demonstrated an interaction between these molecules (Carelli et al., 2019) and HuR is known to be critical to corticogenesis (Kraushar et al., 2014). This inconsistency is likely due to their use of a different isoform of *Pantr2* containing a longer 3 untranslated region. We instead identify a binding site for the RBP Quaking (Qki), presenting a new candidate pathway for *Pantr2* activity. Recent evidence has demonstrated the requirement of an isoform of Qki, Qki-5, for maintenance of multipotency of radial glia (Hayakawa-Yano et al., 2017) through regulation of Ninein alternative splicing (Hayakawa-Yano & Yano, 2019). This interaction has been shown to not only impact fate specification for RGs, but also impact cell cycle dynamics of RGs as Ninein protein is found bound at centrosomes and acts to anchor microtubules during interkinetic nuclear migration (INM) (Shinohara et al., 2013). Loss of Qki-5 also causes RGs to differentiate, raising the possibility that Qki may serve a similar *Pantr2*-dependent role in apical progenitors (Hayakawa-Yano et al., 2017). Qki-5 is also known to regulate N-cadherin and *—*-catenin family members, indicating a potential involvement in regulating Wnt signaling and cell-to-cell adhesion, consistent with our observations of the effect of *Pantr2* in vivo and in vitro (Hayakawa-Yano et al., 2017).

This work lays the groundwork in interpreting the molecular mechanisms used by *Pantr2* and how its absence impacts gene expression, chromatin accessibility and cell cycle dynamics during corticogenesis. We also provide candidate genes that are regulated by *Pantr2* and likely contribute to the *Pantr2* knockout phenotype, including a potential interacting partner in the RBP QKI-5. Finally, we provide in vivo evidence of an altered cell fate trajectory followed by *Pantr2*^-/-^ cells during corticogenesis that results in reduced upper layer fated neurons at E15.5.

## Data Availability

scRNA-seq Pantr2 Neurospheres GSE171636

ATAC-seq Pantr2 Neurospheres GSE210135

snRNA-seq Pantr2 Dorsal Telencephalon GSE210572

## Contributions

Conceptualization: LAG and JJA; Methodology: LAG and JJA; Formal analysis: JJA and ST, Investigation: JJA, ST, TH, BW; Data Curation: JJA; Writing-original draft: JJA, Writing-editing: JJA, LAG, ST, SB; Visualization: JJA and LAG; Supervision: LAG and SB; Funding Acquisition: LAG and JJA; Project Administration: LAG. All authors read and approved the final manuscript.

## Ethical Declarations

All mouse experiments were conducted per the National Institutes of Health Guidance for the Care and Use of Laboratory Animals. All procedures and protocols were approved by the Animal Care and Use Committee of Johns Hopkins University (ACUC protocol number: MO21M54 for mouse cortical neurosphere generation and mouse snRNA-seq library generation).

