## Supplemental Figures for "*Pantr2*, a trans-acting lncRNA, modulates the differentiation potential of neural progenitors in vivo"

### Supplementary Tables

**Supplemental Table 1:** Gene specific primers used for comparative gene expression analysis and RIP analysis in N2a cells
**Supplemental Table 2:** Primers used to barcode samples for ATAC-seq analysis
**Supplemental Table 3:** Primers used for site directed mutagenesis of *Pantr2*
**Supplemental Table 4:** Global differential gene expression list for scRNA-seq in cortical neurospheres
**Supplemental Table 5:** Cell type specific differential gene expression list for scRNA-seq in cortical neurospheres
**Supplemental Table 6:** GO term enrichment analysis for cell type specific differential gene expression list for scRNA-seq in cortical neurospheres
**Supplemental Table 7:** List of unannotated differential peaks found from ATAC-seq in cortical neurospheres
**Supplemental Table 8:** List of annotated differential peaks found from ATAC-seq in cortical neurospheres
**Supplemental Table 9:** GO term enrichment analysis for annotated differential peaks found from ATAC-seq in cortical neurospheres
**Supplemental Table 10:** DNA binding motifs enriched for under differential peaks found from ATAC-seq in cortical neurospheres
**Supplemental Table 11:** GO term enrichment analysis for enriched DNA binding motifs discovered under differential peaks found from ATAC-seq in cortical neurospheres
**Supplemental Table 12:** Cell type specific differential gene expression list for snRNA-seq in E15.5 dorsal telencephalon
**Supplemental Table 13** GO term enrichment analysis for cell type specific differential gene expression list for snRNA-seq in E15.5 dorsal telencephalon

### Supplementary Figures

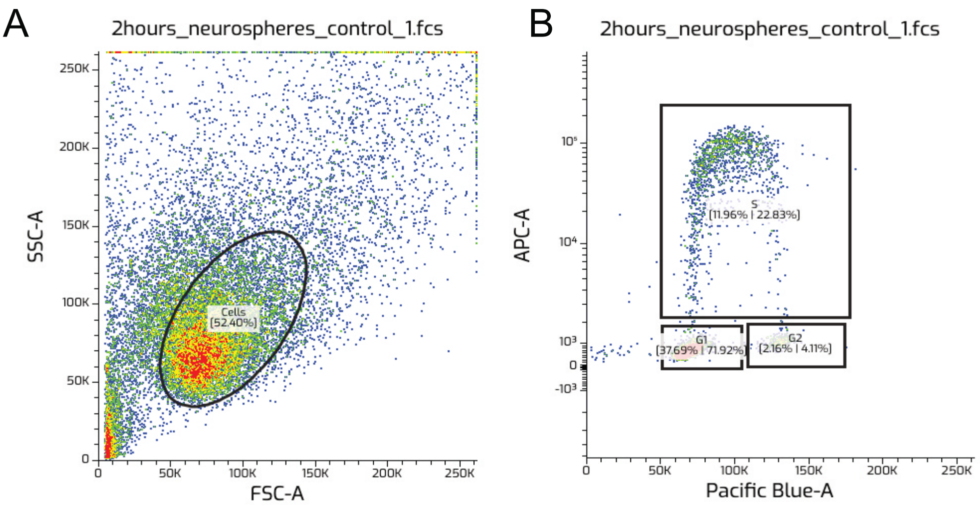


**Figure S1.**
(**A**) Flow cytometry analysis for EdU incorporation assay where whole cells were gated on by SSC-A and FSC-A. A circle gate is drawn around whole, single cells thus filtering out doublets and debris. (**B**) Flow cytometry analysis using the whole cells gated on in panel A where cells are shown by APC-A intensity and Pacific Blue-A. Boxes are drawn around cells found to be in G1, S or G2/M and labeled accordingly.

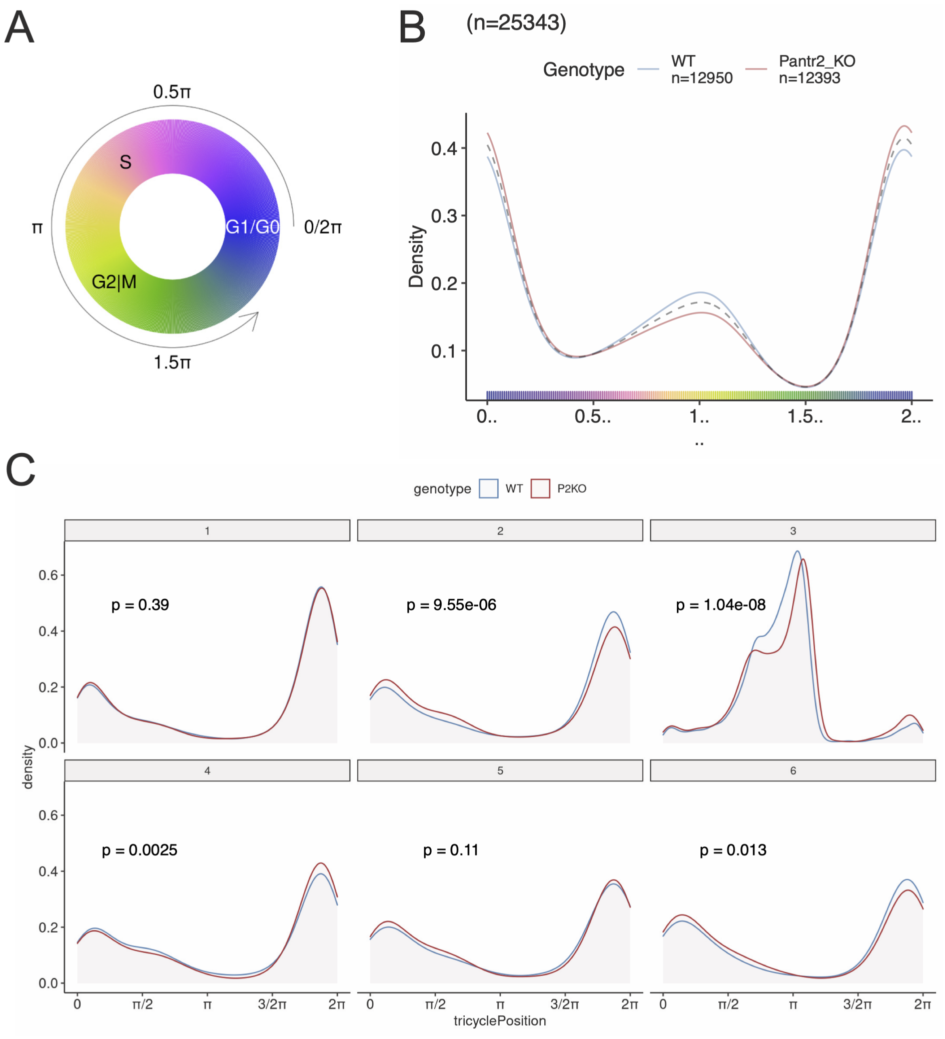


**Figure S2.**
(**A**) Tricycle circular figure legend relating tricycle position from 0-2$\pi$ into cell cycle position G1/G0-S-G2/M. (**B**) Tricycle cell cycle density plot showing the results of the tricycle cell cycle analysis performed on wildtype and *Pantr2*$^{-/-}$cortical neurosphere scRNA-seq data. The density is measured across cell cycle from 0 to 2$\pi$ as shown in the figure legend. (**C**) Tricycle cell cycle density plot showing the results of the tricycle cell cycle analysis performed on wildtype and *Pantr2*$^{-/-}$ E15.5 dorsal telencephalon snRNA-seq data. The density is measured across cell cycle from 0 to 2$\pi$. The p-value is the result of a Wilcoxon Rank Sum Test on tricycle position by genotype for each of the 6 Groups.

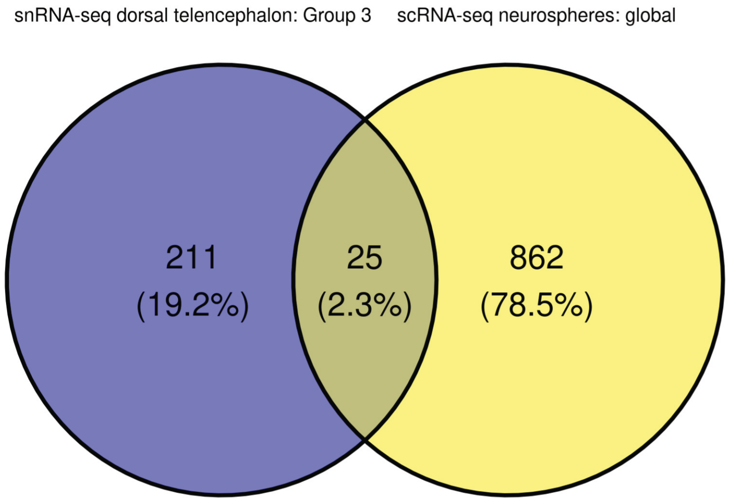


**Figure S3.**
Venn diagram illustrating the overlap in differentially expressed genes between the scRNA-seq performed in cortical neurospheres and the differentially expressed genes found in the snRNA-seq performed in E15.5 dorsal telencephalon.
